# Pigmentation variation in *Drosophila elegans* driven by *cis*-regulatory changes affecting expression of the *ebony* gene

**DOI:** 10.1101/2025.07.28.667336

**Authors:** Anggun Sausan Firdaus, Ateesha Negi, Kai-An You, Erick X. Bayala, Patricia J. Wittkopp, Ben-Yang Liao, Shu-Dan Yeh

## Abstract

Repetitive evolution has offered valuable insights into how similar traits can arise across different populations and species. The dimorphic body color morphs (brown and black) of geographically isolated *Drosophila elegans* Bock & Wheeler, 1972, along with its closely related species *Drosophila gunungcola* Sultana *et al*., 1999, which exhibits black morph, raise intriguing questions about whether these traits have emerged through independent evolution, introgression, or shared ancestral variants. To investigate the genetic and evolutionary origins of these differences, we perform quantitative trait locus (QTL) mapping on recombinant hybrids generated from reciprocal crosses between brown and black morphs *D. elegans*. QTL mapping reveals significant contributions from the X chromosome and Muller element E to intraspecific pigmentation polymorphism, with *ebony* identified as one candidate gene. Analysis of *ebony* expression, including allele-specific expression in F_1_ hybrids, and sequence variation suggests that the intra- and interspecific pigmentation differences within *D. elegans* species subgroup may have evolved independently from the lineage-specific region upstream of *ebony*. By identifying a shared candidate regulatory region associated with both intra- and interspecific variation, our findings suggest that recurrent use of the same genetic region may contribute to the emergence of similar traits across populations and species.

## Introduction

Similar phenotypes within and between species often arise in response to environmental challenges, indicating that ecological conditions often favor similar adaptive solutions. In many cases, such similarities have been linked to divergence in the same genes or pathways, suggesting that certain genetic architectures may be more prone to modification under selection (Bohutinska & Peichel, 2024; Cerca, 2023; Martin & Orgogozo, 2013; Massey & Wittkopp, 2016; Stern, 2013). This raises the intriguing question of how similar phenotypes evolve across populations and species.

Among the phenotypes studied to address this question, pigmentation offers an ideal platform as it is one of the most frequently studied morphological characteristics due to its visibility and ecological relevance that often serve as targets for both natural and sexual selection across taxa (Courtier-Orgogozo et al., 2020; Cuthill et al., 2017). In many *Drosophila* species, pigmentation polymorphisms often appear as a continuous gradient (Brisson et al., 2005; Massey & Wittkopp, 2016; Pool & Aquadro, 2007). These gradients are shaped by ecological pressures and commonly occur across populations living in regions that are geographically proximate (Parkash et al., 2008; Pool & Aquadro, 2007; Telonis-Scott et al., 2011). While pigmentation in many *Drosophila* species follows a continuous gradient, there are cases where geographically isolated populations exhibit distinct body color morphs. The *Drosophila elegans* species subgroup is one particularly notable example of striking dimorphic pigmentation between geographically isolated populations. *Drosophila elegans* populations exhibit two distinct color morphs: brown morph distributed in regions closer to the equator, including Indonesia, the Philippines, and Hong Kong, while a black morph distributed in higher-latitude regions like Taiwan and Okinawa Island (Figure 1a,b). The subgroup also consists of its sister species, *D. gunungcola* (Figure 1c). Notably, *D. gunungcola* is endemic to the southern hemisphere of Sumatra and throughout the Java Islands (Hirai and Kimura, 1997). Despite having partial sympatry with the brown morphs of *D. elegans* at higher elevation, *D. gunungcola* consistently exhibits black morph. This intriguing pattern of pigmentation and distribution raises compelling questions about whether the phenotypic similarities between the black morph of *D. elegans* and *D. gunungcola* are driven by shared genetic variants or independent mutations.

**Figure 1.**
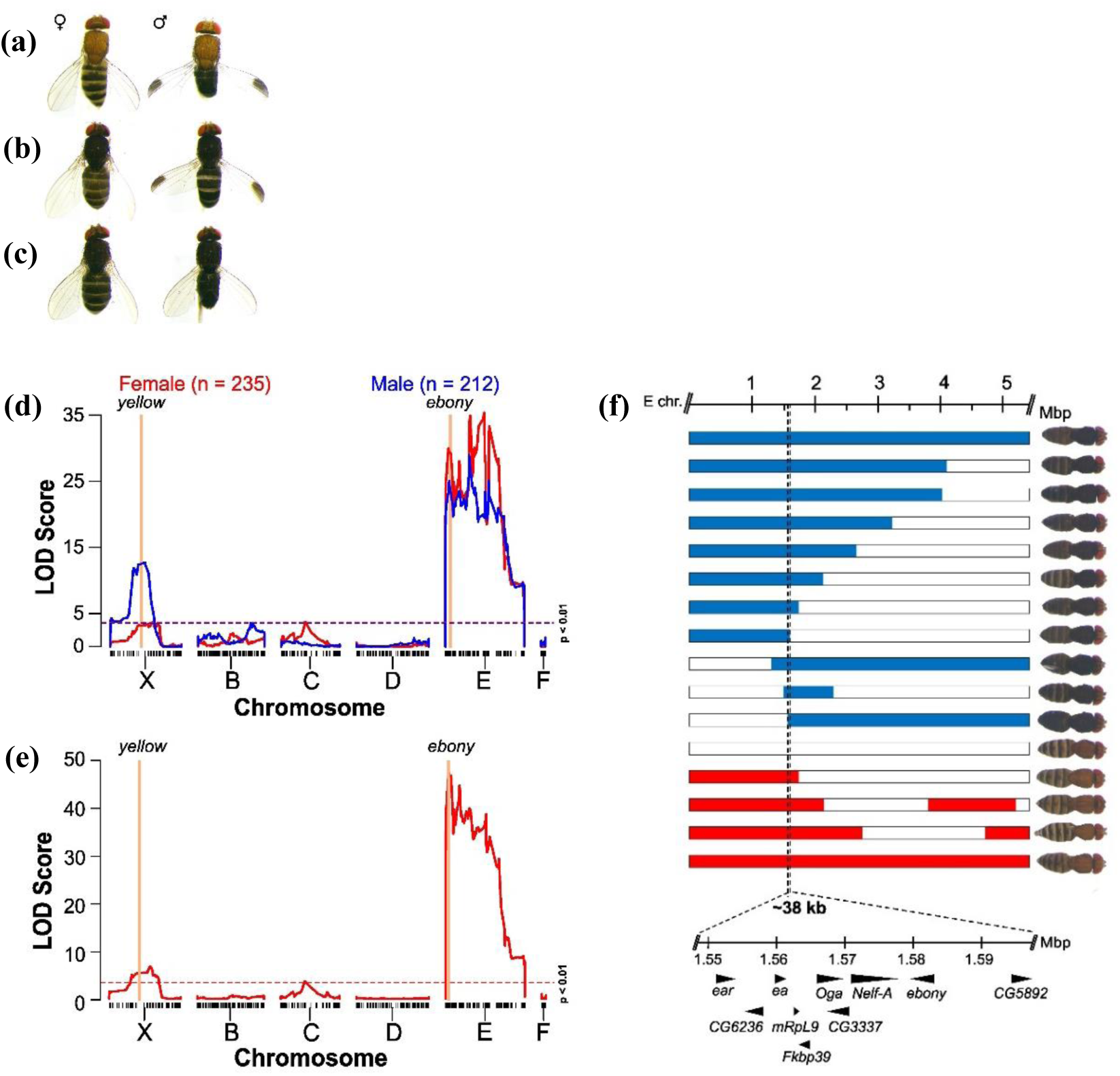
**(a)** Brown morph *D. elegans*. **(b)** Black morph *D. elegans*. **(c)** *D. gunungcola*. **(d)** QTL map for female (red) and male (blue) thorax pigmentation intensity. The y-axis represents LOD score (the red and blue dashed line indicates the p < 0.01 threshold for female and male recombinants, respectively). The x-axis represents the physical map of Muller elements corresponding to *D. melanogaster* chromosome (X = X; B = 2L; C = 2R; D = 3L; E = 3R; and F = 4). Black tick marks on the x-axis represent individual markers along the chromosomes. **(e)** QTL mapping plots for the female abdominal pattern. **(f)** 16 recombinant females containing breakpoints near the thorax pigmentation QTL peaks were aligned to narrow down the candidate loci. Red, blue, and white bars represent homozygous genotype for brown allele, homozygous genotype for black allele, and heterozygous genotype, respectively.

Previous studies have shown that a small set of genes, including *yellow*, *tan*, and *ebony*, are repeatedly implicated in both polymorphic and divergent pigmentation in *Drosophila* (Johnson et al., 2015; Martin & Orgogozo, 2013; Massey & Wittkopp, 2016). For instance, pigmentation polymorphism within *D. americana* and pigmentation divergence between *D. americana* and *D. novamexicana* are both influenced by the genes *ebony* and *tan*, which likely originated from a shared ancestral population (Lamb et al., 2020; Wittkopp et al., 2009). In the *D. elegans* species subgroup, genetic loci linked to the *yellow* and *ebony* genes have been implicated in the pigmentation divergence between the brown morph of *D. elegans* and *D. gunungcola* (Massey et al., 2021). Therefore, we hypothesize that both intra- and interspecific pigmentation variations in the *D. elegans* species subgroup are driven by the same genes. However, whether the intraspecific dimorphism in *D. elegans* and the interspecific divergence involving *D. gunungcola* share common genetic variants remains unknown.

In this study, we explored the genetic basis of dimorphic pigmentation within the *D. elegans* species subgroup, combining QTL mapping, gene expression, and comparative sequence analyses. Our findings identified that differential expression of the pigmentation gene *ebony* likely contributes to both intra- and interspecific variation, potentially through variation in regulatory regions. In particular, a divergent upstream sequence of *ebony* emerged as a candidate region for this dimorphic pigmentation, although the precise causal variants remain unresolved. Together, these findings suggest the potential role of regulatory divergence in shaping similar pigmentation phenotypes within and between species, highlighting the *D. elegans* species subgroup as a valuable model for studying the evolution of dimorphic traits across evolutionary scales.

## Materials and Methods

### Fly strains

*Drosophila elegans* HK (brown morph, originated from Hong Kong, China) and *D. gunungcola* SK (originated from Sumatra Island, Indonesia) strains were provided by John R. True (Stony Brook University) who maintained the stocks established by M. T. Kimura. *Drosophila elegans* TPCC, TPCC12, and TPCC13 strains (black morph) were established from several females collected in New Taipei City, Taiwan. We maintained all strains on a standard cornmeal medium (food recipe is provided by request) under constant light at 23°C in the Department of Life Sciences, National Central University, Taiwan.

The wild-caught flies used for sequencing the *ebony* and mitochondrial DNA were collected from Sumatra and Java Island in 2018 and Taiwan in 2020-2021. The flies were preserved in 75% ethanol upon field collection. Fly collection in Indonesia was carried out under a material transfer agreement with Indonesian collaborators from the Brawijaya University.

### Generating intraspecific recombinants for QTL mapping analysis

Virgin males and females from *D. elegans* HK (brown morph) and TPCC (black morph) strains were categorized 4-6 hours after eclosion and aged for 4-7 days to reach sexual maturity. We started the reciprocal crosses by setting up ten single-pair vials per direction, each containing a male from one strain and a virgin female from another. We allowed each parental pair to copulate and oviposit for 10-14 days, then collected the parental flies for genotyping. We transferred newly eclosed F_1_ recombinants into new vials and allowed them to interbreed for 7-10 days to produce F_2_ recombinants. We maintained the hybrids of each cross pair separately from those of other cross pairs. To promote further recombination, we continued random interbreeding using the same protocol for eight non-overlapping generations.

To map the genetic basis of thorax pigmentation, we phenotyped both male and female recombinants from the 2^nd^ to 8^th^ generations of each reciprocal cross. We visually inspected and classified flies at 7-10 days old. In the 2^nd^ generation, we collected approximately 30 recombinants per color and sex. From the 3^rd^ to 8^th^ generations, we selected the darkest recombinants resembling the TPCC strain and the lightest ones resembling the HK strain (N=12-17 per color and sex).

### Imaging and scoring pigmentation traits

We photographed individual flies to score the pigmentation traits before genotyping. To reduce light reflection interference, we immersed and positioned each fly in 70% ethanol. We captured fly images in RGB mode using a digital camera (MSHOT MS60) connected to a stereoscopic microscope (Leica MZ 125), applying identical optical parameters for all samples. We quantified thorax pigmentation intensity using ImageJ software (version 1.52a; Wayne Rasband, National Institutes of Health, USA; http://imagej.nih.gov/ij). We manually selected six regions of interest (ROIs), modified from (Miyagi et al., 2015), including the thoracic trident, thoracic scutellum, and areas outside the trident. For each ROI, we recorded the mean gray value. To align scores with a scale ranging from 0 (white) to 255 (black), we substracted each score from 255 (Gibert et al., 2016). We averaged the six scores to obtain the final pigmentation intensity score for each sample. Intensity values are reported in arbitrary units (A.U.).

For the female abdominal banding pattern, we measured the width of the dark band and the total width of the 4^th^ tergite. We calculated the percentage of coverage coverage by dividing the dark band width by the total tergite width and multiplying the result by 100.

### Library preparation, genome assembly, and marker generation

A total of 469 flies, including 226 male and 243 female recombinants, were individually genotyped using Multiplexed Shotgun Genotyping (MSG) (Andolfatto et al., 2011) with 473 barcoded adaptors, following the protocol described in (Cande et al., 2012). We extracted genomic DNA and prepared sequencing libraries using the methods explained in (Massey et al., 2020). For read mapping, we used the *D. elegans HK* reference genome (D.elegans_HK_3.0; accession number: GCA_011057505.1). This genome reference has been assembled to chromosomal-level which facilitates more precise mapping to specific chromosomal locations. We sequenced the TPCC strain genome at 50x coverage allowing a confident variant calling between HK and TPCC strain. In total, we identified 3198 informative markers between the two strains for QTL mapping. The markers, thorax and female abdominal phenotypes, and script used for QTL mapping are available on [repository to be determined].

### QTL mapping analysis and gene annotation

We conducted QTL mapping analysis using the R/qtl package (Broman et al., 2003) in R for Windows version 4.4.0 (R Core Team, 2024). We inferred ancestry states across chromosomal regions, recombinant breakpoints, and genotyping status using a Hidden Markov Model developed for MSG data, and imported the results into the R/qtl package using a custom script (https://github.com/dstern/read_cross_msg) (Andolfatto et al., 2011). We performed genome scans using Haley-Knott regression and the “scanone” function of a single QTL model (Haley & Knott, 1992). Due to the low sequencing resolution, we excluded 13 males and 8 females from the analysis, resulting in a final dataset of 212 males and 235 females from the 2^nd^ to 8^th^ generations. We determined the LOD significance threshold using 1,000 permutation replicates at a 1% significance level (α = 0.01). We extracted the chromosomal regions spanning the QTL interval (∼38 kb on Muller element E) for thorax and abdominal pattern from the assembled *D. elegans* HK genome (accession no: CM021661.1). Gene annotation followed the method described in Massey et al. (2020).

### Analyzing epistasis interactions between QTL-X and ebony

To investigate potential epistatic interactions between QTL on the X chromosome and Muller element E, we compared thorax pigmentation intensity (in both males and females) and female abdominal banding patterns across different genotypic combinations. The QTL-X genotypes were identified at positions with the highest LOD scores from our QTL mapping (Table 1; Male thorax pigmentation intensity: 12,486,125 bp; Female thorax pigmentation intensity: 15,476,093 bp; Female abdominal dark band width: 14,725,205 bp). For Muller element E, we focused on a region previously narrowed down (position: 1,582,436 bp, located near *ebony*).

**Table 1.** QTLs detected for thorax pigmentation intensity and abdominal banding pattern.

| Traits | Sex | Chromosome | QTL interval (bp)<sup>a</sup> | QTL peak (bp) | LOD |
| --- | --- | --- | --- | --- | --- |
| Thorax pigmentation intensity | Female | X | 7,445,655-18,244,170 | 15,476,093 | 3.71 |
| Thorax pigmentation intensity | Male | X | 7,955,254-14,476,391 | 12,486,125 | 12.72 |
| Thorax pigmentation intensity | Female | E | 8,622,548-14,156,589 | 14,038,360 | 35.38 |
| Thorax pigmentation intensity | Male | E | 8,422,980-8,972,186 | 8,487,591 | 29.01 |
| A4-tergite dark band width | Female | X | 7,955,254-15,699,310 | 14,725,205 | 6.77 |
| A4-tergite dark band width | Female | E | 600,044-2,080,945 | 1,069,099 | 47.11 |

<sup>a</sup>1.8-LOD drop support interval

### RNA Sample Preparation and RT-qPCR

We assessed the relative gene expression of *ebony* between brown (HK strain) and black (TPCC12 strain) morphs of *D. elegans* using RT-qPCR. We used a different black morph strain collected from the same locality as TPCC because the TPCC strain originally used for QTL mapping perished several months after experiments. We also included *D. gunungcola* (SK strain) to test whether its *ebony* expression was similar to black morph *D. elegans*, as both share similar body pigmentation.

We collected male and female flies within 0 to 2 hours after eclosion. After removing the wings, we stored the bodies in RNAlater^TM^ Solution (Thermo Fisher Scientific) for dissection. We dissected flies into three body parts: head, thorax epidermis (including scutellum and pleura), and the dorsal side of abdominal epidermis. For each body part, sex, and strain, we prepared three biological replicates (∼40 flies per replicate). We extracted Total RNA using the Quick-RNA^TM^ Miniprep Kit (Zymo Research, USA) following the manufacturer’s instructions and quantified the RNA using Nanodrop spectrophotometer. First-strand cDNA was synthesized from ∼200 ng of total RNA using the MMLV Reverse Transcription Kit (PROTECH).

We performed RT-qPCR using a Bio-Rad CFX96 system in 20-µl reaction containing SYBR Green Supermix (BioRad, USA) and 1 µl of cDNA per reaction. Primer sequences used for *ebony*, *Actin42A*, and *TBP* are listed in Table S5. Relative expression of *ebony* was calculated using the Pfaffl method (Pfaffl, 2001), with *Actin42A* and *TBP* as reference genes. Whole-body RNA from *D. elegans* HK was used as the control sample.

### RNA and DNA extraction for allele-specific expression (ASE) analysis

To investigate cis-regulatory divergence of the *ebony* gene between pigmentation morphs in *Drosophila elegans*, we performed allele-specific expression (ASE) analysis using a Sanger sequencing. We used genomic DNA from the parental strains, HK strain (brown morph) and TPCC13 strain (black morph), as reference samples for normalization and baseline correction. We pooled five adult females and extracted DNA using the GeneAid gSYNC DNA Extraction Kit. We constructed a standard curve for normalization by mixing parental gDNA samples at the brown:black ratios of 1:10, 1:5, 1:4, 1:2, 1:1.5, 1:1, 1.5:1, 2:1, 4:1, 5:1.

We crossed 5 HK females with 5 TPCC13 males to obtain F_1_ recombinants. To prepare the total RNA from F_1_ recombinants, we dissected females within 0-2 hours post-eclosion to isolate the head, thorax epidermis, and abdominal epidermis. For each tissue type, we prepared three biological replicates, each consisting of 20 pooled individuals. We extracted total RNA using the Gene-Spin Total RNA Purification Kit (PROTECH), following the manufacturer’s protocol. We reverse-transcribed approximately 200 ng of RNA per replicate into cDNA using the MMLV Reverse Transcription Kit (PROTECH).

We PCR-amplified the target region (∼800 bp of *ebony* exon 2) from gDNA and cDNA samples using e_F9 and e_R8 primers (Table S5). PCR conditions were: initial denaturation at 95°C for 2 min, followed by 35 cycles of 95°C for 30 s, 59°C for 1 min, and 72°C for 20 s, with a final extension at 72°C for 5 min. We sent the samples for Sanger sequencing using the forward primer (PB_e_F9).

### Allele-Specific Expression Quantification and Normalization

To quantify allele-specific expression, we extracted peak height values from sequencing chromatograms (.ab1) using QSVanalyser (Carr et al., 2009) for each F_1_ biological replicate. We used gDNA from each parental strain (HK accession number: PV939603; TPCC13 accession number: PV939604) to identify informative reference SNPs that distinguish the brown and black alleles of *D. elegans*. We focused on a single reliable marker SNP (located at 489 bp), which was fixed between the brown and black parental strains and showed consistent allelic imbalance across all tissues and biological replicates. We excluded other candidate SNPs from the analysis because they were not fixed (i.e., both alleles were present in at least one parental strain, as indicated by their chromatograms).

We corrected the final peak heights using baseline signal values following the method of Carr et al. (2009). For each replicate, we calculated the allelic expression ratio by dividing the final peak height of the brown allele by the black allele. We normalized these ratios using the standard curve generated from mixed parental gDNA samples with known brown:black ratios. This normalization accounted for technical variation in peak heights and allowed us to more accurately estimate the relative expression of the two alleles.

### Hybridization Chain Reaction (HCR)

We performed Hybridization Chain Reaction (HCR) *in situ* staining to detect mRNA expression of *ebony* and *yellow* in *Drosophila elegans* black (TP) and brown (HK) morphs, following the protocol described in (Bayala et al., 2024). DAPI was included to visualize DNA. The same custom hairpins targeting *yellow* and *ebony* described in this prior work were also used in the current study. To assess spatiotemporal gene expression, we stained whole-body samples at two developmental stages: 85-95 hours after puparium formation (hAPF) and 0-2 hours post eclosion. We did not select for specific sexes, as the objective was to capture gene expression at carefully matched developmental stages. To further examine spatiotemporal expression in the abdominal epidermis during pupal development, we additionally stained dissected pupal abdomens at 65-75 hAPF and 75-85 hAPF. After staining, tissues were mounted on slides and imaged using a Leica SP5 laser scanning confocal. Red, green, and blue channel levels were adjusted uniformly across all image panels.

### Collection of wild D. elegans in Indonesia and Taiwan

We collected samples in Indonesia for over three months in 2018 across Sumatra and Java. In Taiwan, we conducted sample collection in 2020 and 2021. Flies were collected using an aspirator and transferred into vials containing standard fly media or Instant Drosophila medium (agar, glucose, cornmeal, and yeast extract). Some flies were immediately preserved in ethanol 70%, and the rest of the caught flies were maintained and reared in the laboratory to establish isofemale or wild-isolated strains. Fly collection in Indonesia was carried out under a material transfer agreement with Indonesian collaborators from the University of Brawijaya.

### Haplotype sequence analysis of ebony region

We extracted genomic DNA from single adult male flies using Gsync^TM^ DNA Extraction Kit, following the manufacturer’s instruction. We analyzed a total of 20 individuals: 9 *D. elegans* and 6 *D. gunungcola* from Indonesia (Java and Sumatra), and 5 *D. elegans* from Taiwan (Table S2). We used six pairs of primers targeting partially overlapping ∼3 kb amplicons to amplify ∼16 kb of the *ebony* locus (Table S5). The amplified region spanned the entire *ebony* gene (including 5’ and 3’ intergenic region, introns, and exons).

We assessed PCR product quality using gel electrophoresis and checked the quantity using a Qubit® fluorometer (ThermoFisher Scientific) with High Sensitivity (HS) dsDNA assay kits and a NanoDrop spectrophotometer. We sent the amplicons for sequencing using the PacBio Single-Molecule Real-Time (SMRT). We evaluated the quality of the amplicon-specific consensus sequences using SequelTools program/SequelQC v1.3.0. (Hufnagel et al., 2020). For each sample, six read clusters corresponding to the six primer pairs were phased into haplotypes. We mapped the reads to the *D. elegans* reference genome (D.elegans_HK_3.0; accession number: GCA_011057505.1) using the pbmm2 program (https://github.com/PacificBiosciences/pbbioconda). We visualized variants from each cluster in IGV, carefully selected for each respective amplicon by aligning it to reference using NUCmer from the MUMmer suite version 3.1, and stitched into a complete haplotype (Kurtz et al., 2004).

We aligned DNA sequences using ClustalW in MEGA-X v10.2.5 (Kumar et al., 2018). To examine patterns of genetic divergence, we performed a 100-bp sliding-window analysis of net nucleotide divergence (*Da*) across the ∼16 kb region spanning *ebony* regions and its neighboring genes using DnaSP v6.12.03 (Rozas et al., 2017). Sample information, including sample ID, accession numbers, and localities is described in Table S2.

### Phylogenetic tree reconstruction

We reconstructed phylogenetic trees using Bayesian Inference implemented in MrBayes v3.2.7a (Ronquist et al., 2012). We analyzed three datasets separately: the complete coding region of *ebony*, 400 bp of the 5’ intergenic region, and mitochondrial DNA (*ND1*: NADH dehydrogenase subunit 1). For the mitochondrial dataset, we used a different set of samples due to limited availability. Genomic DNA was extracted using Gsync^TM^ DNA Extraction Kit, following the manufacturer’s protocol. We amplified the *ND1* region using primers (mtF9 and mtR14) listed in Table S5. The samples used for each analysis are detailed in Table S2.

For all datasets, we used *D. fuyamai* as an outgroup. For the *ebony* region, we extracted the coding sequence along with upstream intergenic regions from the genome assembly of *D. fuyamai* (accession number: JAECXW010000874.1). For the mitochondrial dataset, we used *ND1* sequence from *D. fuyamai* (accession number: HQ631806). Due to poor sequence conservation, the alignment quality of the 400-bp upstream region of *ebony* with the outgroup was not optimal. Despite this limitation, we included it in the phylogenetic reconstruction to explore potential evolutionary relationships.

To determine the best-fitting nucleotide substitution model for each dataset, we used jModelTest 2.1.10 v20160303 (Darriba et al., 2012) and selected models based on the Bayesian Information Criterion (BIC). Convergence was confirmed by examining the average standard deviation of split frequencies (<0.01). We discarded the first 25% of sampled trees as burn-in and summarized the remaining trees using a 50% majority-rule consensus to construct the final phylogeny.

### Statistical analysis

We conducted all statistical analyses in R for Windows v4.4.0 (R Core Team, 2024), unless otherwise specified. Specific statistical analysis approaches varied by analysis and are described in detail in the relevant method subsections (e.g., QTL mapping, epistasis analysis). For RT-qPCR, we used one-way ANOVA, followed by Tukey’s HSD test for post hoc pairwise comparisons. For allele-specific expression, we applied one-sample t-tests on log-transformed allele ratios to evaluate significant deviations from equal expression (1:1 ratio) between alleles. We used two-way ANOVAs for epistasis analyses.

## Results

### The X chromosome and Muller element E contribute to intraspecific pigmentation dimorphism

To examine the inheritance of intraspecific pigmentation variation, we performed reciprocal crosses between two strains of *D. elegans*: HK (brown morph) and TPCC (black morph). The F1 flies from both crosses exhibited intermediate body color (Figure S1a). We then generated sibling crosses from the F1 hybrids for eight generations. We focused our analysis on the thorax pigmentation polymorphism in both male and female recombinants. The resulting recombinants displayed a spectrum of thorax pigmentation intensity.

To investigate the candidate loci linked to the thorax pigmentation variation, we performed QTL mapping on 235 females and 212 males from F2 to F8 recombinants produced through reciprocal crosses using individuals with the body color at two ends of the phenotypic distribution (Figure S1b). There were no significant differences in thorax pigmentation intensity between male and female recombinants within the brown morph or within the black morph, indicating that this trait is not sexually dimorphic (Figure S2).

Our QTL mapping analyses revealed an X-linked QTL that was highly significant in the male mapping dataset, but only marginally significant in the female mapping dataset (Figure 1d; Table 1). This difference in significance presumably results from differences in mapping power for X-linked loci in the two sexes: males are hemizygous for X-linked genes, carrying only the black or brown morph allele, whereas females include individuals that are homozygous for the black morph allele, homozygous for the brown morph allele, or heterozygous for both. The entire Muller element E (corresponding to the 3R chromosome in *D. melanogaster*) exhibited strong associations with multiple peaks in both male and female mapping datasets (Figure 1d; Table 1). The broad QTL signals across Muller element E may be attributed to: 1) multiple QTLs of large effect; 2) low resolution from the relatively small number of recombinant hybrids; or 3) a low recombination rate on Muller element E. The latter scenario is less likely as crossover frequencies were consistent among chromosomes (Figure S3).

We also observed variations in female abdominal banding patterns (Figure 1e). To compare the genetic basis of thorax pigmentation intensity and abdominal banding pattern variation, we measured the width of dark bands on the 4th tergite of 235 female F2 to F8 recombinants. We detected a significant positive correlation between thorax pigmentation intensity and the width of dark band on the A4 (Spearman’s correlation: *R*=0.76, *P* < 0.001) (Figure S4), suggesting that darker thoraxes are associated with wider abdominal dark bands. Our QTL mapping analyses identified a highly significant QTL on Muller element E associated with variation in the abdominal banding pattern. A QTL with lower significance on the X chromosome also showed genetic association with female abdominal banding pattern (Figure 1e; Table 1). This was unexpected, as thorax pigmentation intensity and spatial banding pattern in the abdomen are typically regulated by different genetic mechanisms (Massey & Wittkopp, 2016; Rebeiz et al., 2009; Takahashi, 2013). Our results suggest these traits may share a genetic basis, potentially controlled by overlapping or closely linked regulatory elements.

*Candidate loci within QTLs and genetic interaction between X chromosome and Muller element E* Muller element E contains multiple genes known to be associated with pigmentation in *Drosophila*, including *ebony*, *Abdominal-B (abd-b)*, *Dop1R1*, *Dop1R2*, and *stripe (sr)*. The QTL signal spans a large portion of Muller element E, suggesting that one or more known or previously uncharacterized genes may contribute to pigmentation differences. To refine the candidate region on Muller element E, we examined recombinant genotypes and phenotypes to identify the recombination breakpoints flanking the thorax QTLs. This allowed us to narrow down only one candidate region to ∼38 kb in females, containing 10 genes in the region acting like a genetic switch controlling brown or black thorax pigmentation (Figure 1f). In males, we refined the region to ∼891 kb, which includes the region identified in females (Figure S5). Hybrids homozygous for the black loci in this region exhibited black thorax pigmentation, whereas those with heterozygous or homozygous brown loci displayed brown thorax (Figure 1f, 2a). One gene in this region is *ebony* (*e*) which encodes the enzyme N-ß-alanyl-dopamine synthetase required for the formation of yellow sclerotin (Wittkopp et al., 2002). *ebony* has been previously reported to contribute in pigmentation variation both within and between *Drosophila* species (Johnson et al., 2015; Massey & Wittkopp, 2016; Wittkopp et al., 2009), making it a strong candidate gene responsible for pigmentation variation in *D. elegans*.

We also attempted to refine the candidate region on the X chromosome associated with thorax pigmentation. However, we were unable to identify a specific region on the X chromosome, as individuals carrying homozygous black, heterozygous, or homozygous brown loci across the entire X chromosome could exhibit either brown or black thorax pigmentation (Figure S6). To test for possible genetic interactions, we further performed a two-way ANOVA between the QTL identified on the X chromosome (hereafter referred to as QTL-X) and the *ebony* locus on Muller element E. Our analysis revealed tissue- and sex-specific genetic interactions between *ebony* and QTL-X, using the marker with highest LOD score (Figure 2b; Two-way ANOVA; Male thorax pigmentation intensity: *F_2,206_* = 2.79, *P* = 0.064; Female thorax pigmentation intensity: *F_4,226_* = 2.63, *P* = 0.035; Female abdominal dark band width: *F_4,226_* = 2.21; *P* = 0.069). We observed that brown *ebony* allele was dominant to black *ebony* allele regardless of QTL-X allele for thorax pigmentation. While male recombinants showed a marginally significant interaction, female thorax pigmentation showed weak epistatic interaction. These results suggest that while *ebony* is the primary driver of pigmentation, QTL-X likely contributes through additive effects and weak epistatic interactions in a tissue- and sex-specific manner.

**Figure 2.**
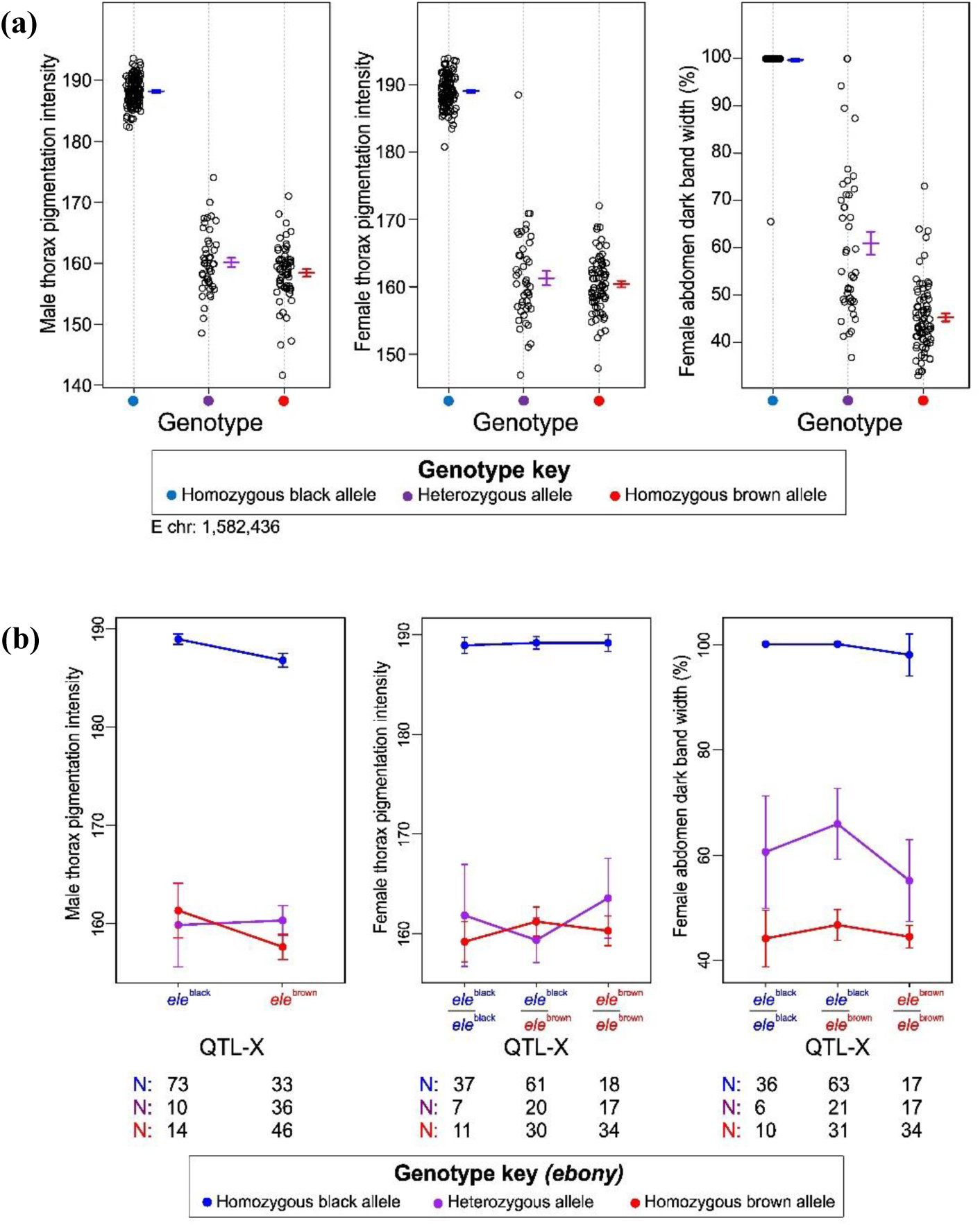
**(a)** Effect plots of the genotype against QTL peak (marker: 1,582,436 near the *ebony* locus) for male thorax pigmentation intensity, female thorax pigmentation intensity, and female abdominal pattern. Dots represent individual replicates. **(b)** Genetic interactions between QTL on the X chromosome (QTL-X) and *ebony* on Muller element E. Error bars represent 95% confidence intervals for the phenotypic values of each genotype. Sample sizes (N) for flies with each genotype (blue for homozygous *D. elegans* black allele, purple for heterozygous allele, and red for homozygous *D. elegans* brown allele) are provided below each plot for the respective traits. Higher score means darker pigmentation intensity.

### Variation in ebony expression is correlated with Drosophila elegans pigmentation differences

Given the pleiotropic role of *ebony* in *Drosophila* (Borycz et al., 2002; Massey et al., 2019; Takahashi, 2013), we hypothesized that genetic variation in *ebony* might regulate its expression levels rather than altering its protein function, contributing to both intra- and interspecific pigmentation differences in the *D. elegans* species subgroup. To test this, we performed RT-qPCR on brown morph *D. elegans*, black morph *D. elegans*, and black morph *D. gunungcola*. The inclusion of *D. gunungcola* allowed for a comparison between closely related species with the same body color morphs. We collected male and female samples within 0 to 2 hours post-eclosion, and focused on the head, thorax epidermis, and abdominal epidermis to capture tissue-specific expression patterns. We observed consistent *ebony* expression patterns across tissues and sexes. Generally, the brown morph showed significantly higher *ebony* expression than the black morphs (Figure 3a). Notably, the thorax epidermis showed especially strong differences in *ebony* expression between brown and black morphs in both male and female datasets.

**Figure 3.**
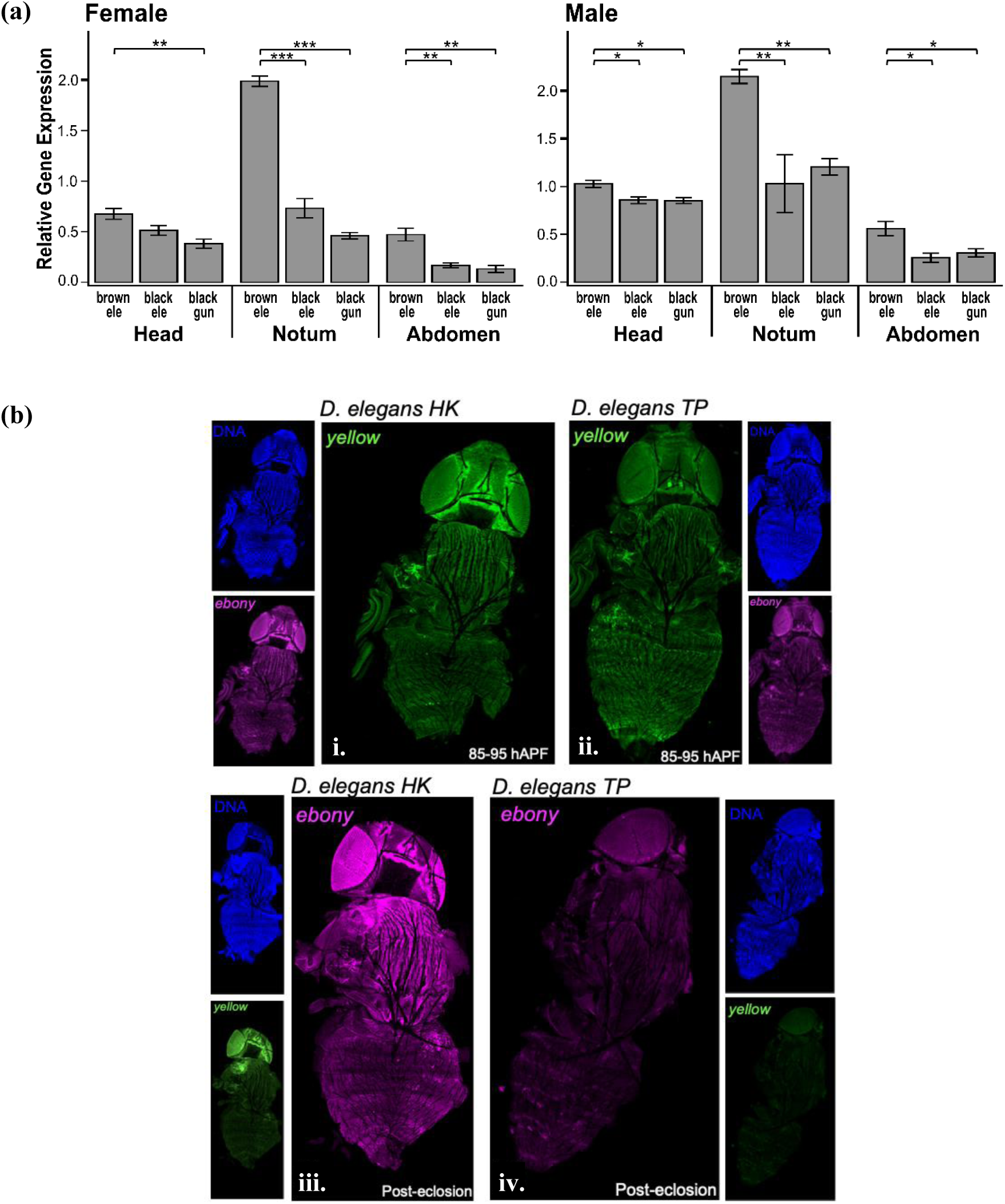
**(a)** Relative *ebony* gene expression among different body morphs. Newly emerged flies were dissected into three parts: head, thorax epidermis, and dorsal abdominal epidermis. One-way ANOVA followed by *post hoc* Tukey HSD were used to assess significant differences among group. Significant *P value*: ‘*’ ≤ 0.05; ‘**’ ≤ 0.01; ‘***’ ≤ 0.001. **(b)** Spatiotemporal expression of *ebony* and *yellow* in black and brown morphs of *D. elegans*. The epidermal tissues from brown morph (*D. elegans* HK) and black morph (*D. elegans* TP) were stained using the Hybridization Chain Reaction (HCR) at two developmental stages: (i; ii) 85–95 hours after puparium formation (hAPF); and (iii; iv) 0–2 hours post-eclosion. For each sample, three panels are shown: DNA (DAPI, blue), *ebony* mRNA (magenta), and *yellow* mRNA (green). All samples were processed and imaged under the same conditions; variation in DAPI signal may reflect staining efficiency or nuclear accessibility.

To test whether these *ebony* expression differences stem from *cis*-regulatory divergence, we analyzed expression of the *ebony^HK^* (the brown morph allele) and *ebony^TP^* (the black morph allele) in F_1_ hybrids after eclosion. The *ebony^HK^* consistently showed higher expression than the *ebony^TP^* across all tissues (one-sample *t*-test, *P* < 0.05, Table S1), indicating that *cis*-regulatory divergence does indeed contribute to the difference in expression level seen between these two morphs.

We further investigated the spatial distribution of *ebony* and *yellow*, a pigmentation gene within QTL-X peak, mRNA expression of black and brown morphs of *D. elegans* using the Hybridization Chain Reaction (HCR) method. The brown morph displayed stronger *ebony* expression than the black morph at 65-75 hours after puparium formation (hAPF) (Figure S7) and this difference become more pronounced at later developmental stages (75-85 hAPF, 85-95 hAPF, and post-eclosion), aligning with RT-qPCR findings (Figure 3b; Figure S7). In contrast, *yellow* expression was detected in the epidermis in both morphs, though the signal appeared slightly stronger before eclosion in the black morph and after eclosion in the brown morph. These findings correlate with the effects observed for QTL-X and QTL-E, with larger differences in *ebony* expression than *yellow* expressing between the black and brown morphs. Together, our results from three kinds of gene expression analyses support that *cis*-regulatory differences in *ebony* underlie intraspecific pigmentation dimorphism in *D. elegans*.

### 5’ intergenic region of ebony displayed high nucleotide variation that correlates with dimorphic pigmentation differences at both intra- and interspecific levels

To explore natural genetic variation within the *ebony* locus that may be associated with pigmentation dimorphism, we sequenced a ∼16 kb *ebony* region (including the 5’ and 3’ intergenic, exons, and introns) from wild-caught *D. elegans* and *D. gunungcola* from Indonesia and Taiwan (Table S2). We first investigated protein sequence and found no fixed amino acid differences between brown and black *D. elegans* morphs. In contrast, we identified seven amino acid residues diverged between two species (Table S3). Five of these residues are located within the adenylation domain of the Ebony protein, hypothetically activate ß-alanine to aminoacyladenylate for further conjugating on to dopamine (Izore et al., 2019; Richardt et al., 2003). The functional significance of these five residues, however, have not been experimentally tested. The remaining two sites are located in the C-terminal region (Izore et al., 2019). Based on the *D. melanogaster* Ebony structure, none of these residues is at critical sites for substrate binding, orientation, or catalysis, suggesting they are unlikely to influence Ebony’s enzymatic function.

Instead of protein sequence differences, *cis*-regulatory changes in the *ebony* locus may result in pigmentation differences. To explore lineage-specific versus shared genetic patterns linked to pigmentation, we performed a sliding-window analysis of net divergence (*Da*) using the ∼16 kb *ebony* across three pairwise comparisons-between brown and black *D. elegans* as well as *D. gunungcola*. Between-species comparisons exhibit higher nucleotide divergence than within-species comparisons generally as expected, except in the region of 3635 bp to 4035 bp from the end of upstream gene coding sequence. This 400-bp sequence in the 5’ intergenic region of *ebony* shows high divergence between brown and black morphs for both within and between species comparisons, but not between black morph *D. elegans* vs. *D. gunungcola* (Figure 4; Figure S8). The relatively low divergence between black morph *D. elegans* and *D. gunungcola* implies that natural selection may have acted on this region in the black morph to reflect similar evolutionary pressures in both species or shared ancestral genetic variants.

**Figure 4.**
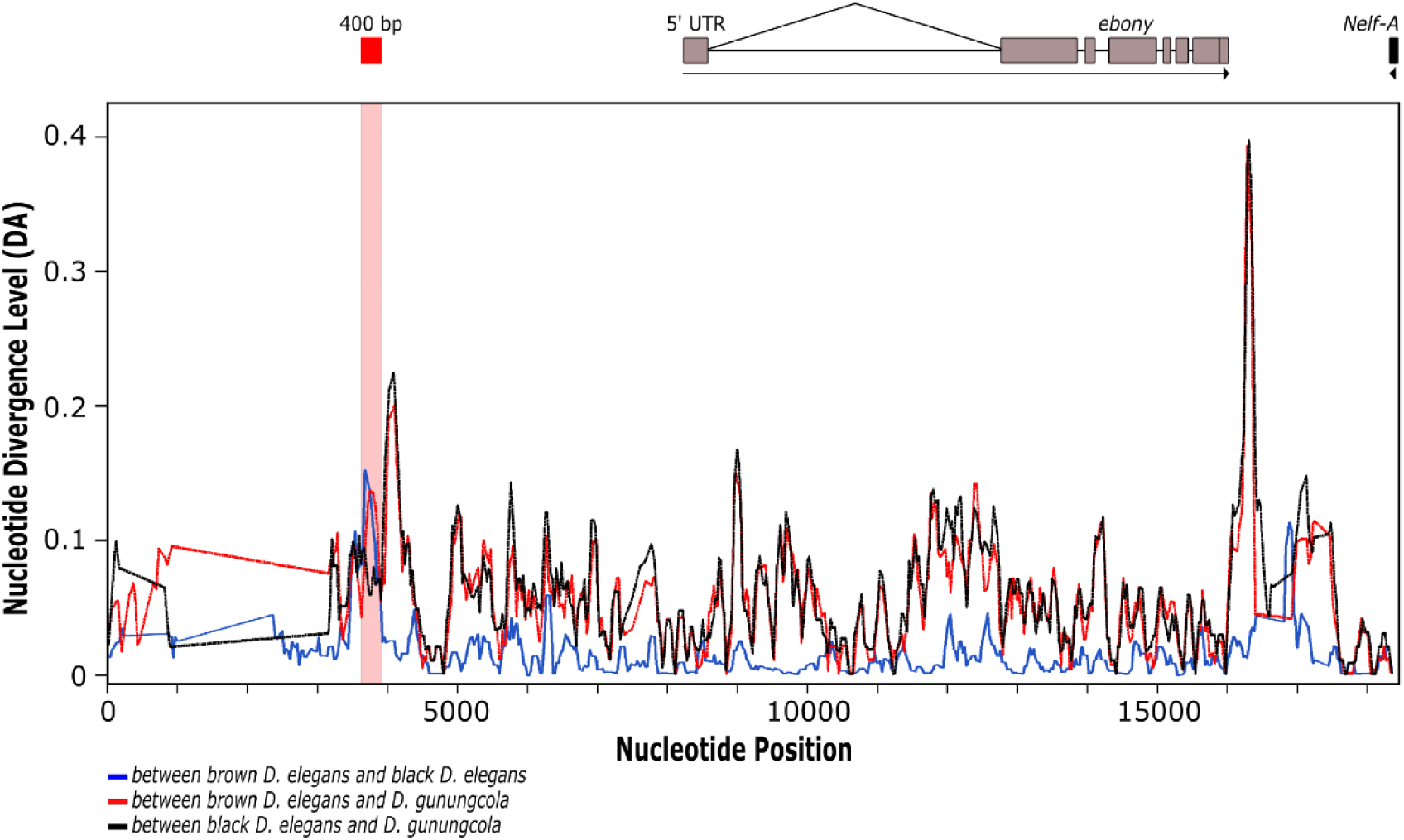
Sliding window analysis of net divergence levels (DA) between *D. gunungcola* and *D. elegans*. Three pairwise comparisons were performed: brown *D. elegans* vs. black *D. elegans* (blue), brown *D. elegans* vs. *D. gunungcola* (red), and black *D. elegans* vs. *D. gunungcola* (black). Grey, black, and red (400 bp) horizontal bars at the top of the figure indicate the transcribed regions of the *ebony* gene, nearby gene, and *ebony* region with high divergence levels. Window size was set to 100 bp with step size 25 bp.

### Phylogenetic patterns of dimorphic pigmentation in D. elegans species subgroup

To investigate the evolutionary histories underlying dimorphic pigmentation in *D. elegans* species subgroup, we reconstructed phylogenetic trees of wild-caught samples using three genomic regions: the 400-bp candidate region in the 5’ intergenic region of *ebony*, the coding sequence (CDS) of *ebony*, and mitochondrial DNA (*ND1*) (Table S2). Examining these regions separately is important as each may reflect distinct evolutionary trajectories and selective pressures.

The tree based on *ebony* coding sequences formed two major clades: *D. gunungcola* clade and *D. elegans* clade, within which the black morph is nested in the brown morph clade (Figure 5a). In contrast, the tree based on the 400-bp 5’ intergenic region of *ebony* formed three major clades: *D. gunungcola* clade, brown morph *D. elegans* clade, and black morph *D. elegans* clade (Figure 5b). On the other hand, the mitochondrial DNA tree did not separate black and brown *D. elegans* into distinct clades (Figure 5c). While the phylogenetic tree based on the 400-bp candidate region revealed distinct clades for the brown and black morphs of *D. elegans*, this pattern contradicts our hypothesis that the black morphs of *D. elegans* and *D. gunungcola* share ancestral genetic variants for body melanization.

**Figure 5.**
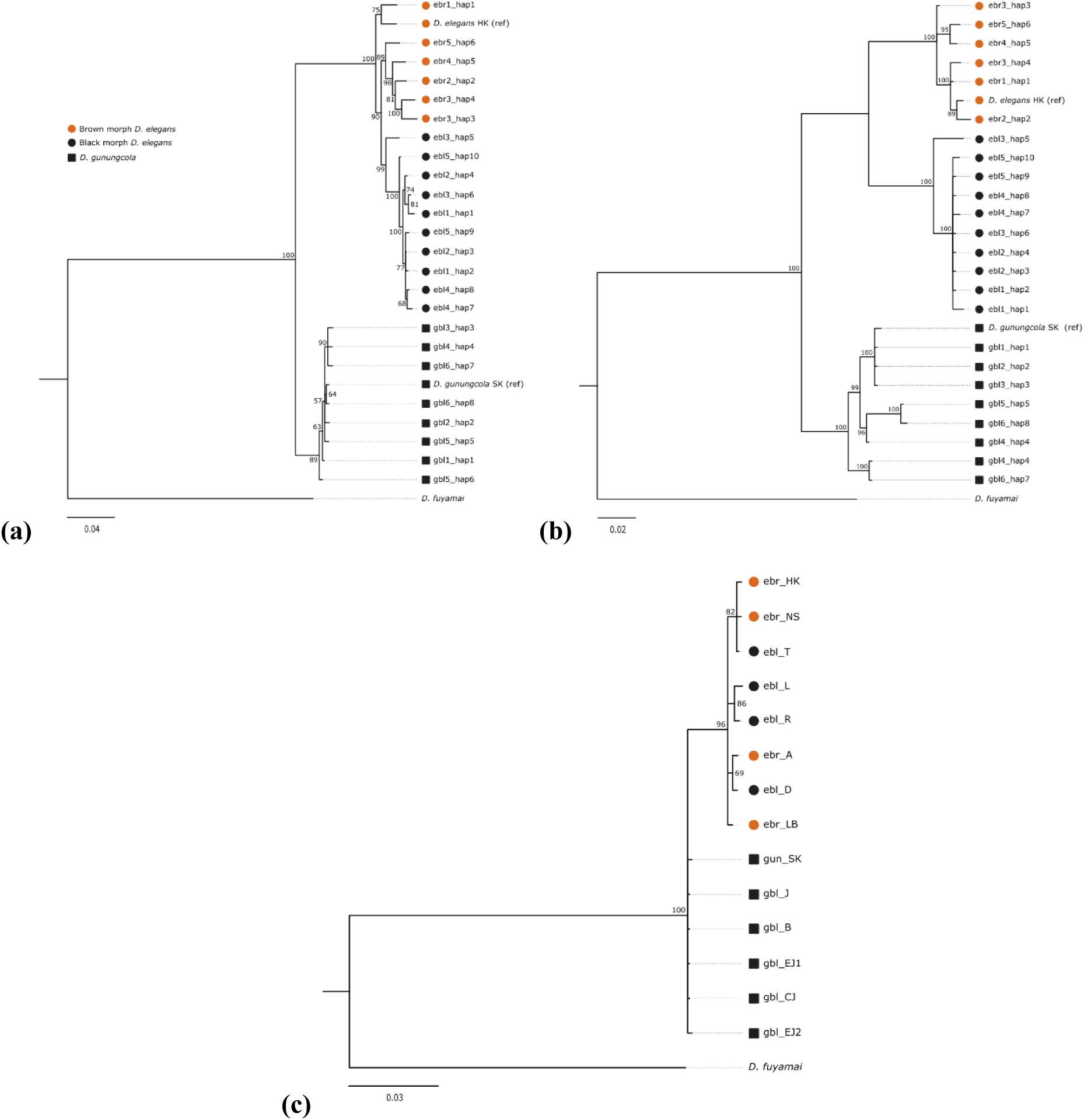
Phylogenetic trees reconstructed using Bayesian Inference based on **(a)** the *ebony* coding sequence, **(b)** the 5’ intergenic region of *ebony*, and **(c)** mitochondrial DNA (*ND1*). Numbers on the branches represent posterior probabilities from Bayesian analysis, indicating statistical support for each node.

If dimorphic pigmentation is resulted from shared genetic variants, we would expect a substantially higher number of shared polymorphisms exclusive to black *D. elegans* and *D. gunungcola*. We analyzed the sequence polymorphisms within the 400-bp candidate region and identified 18 diverged polymorphisms (16 nucleotide substitutions and 2 indels) between species, 12 shared nucleotide substitutions exclusive to black morph *D. elegans* and *D. gunungcola,* and 11 shared polymorphisms (10 nucleotide substitutions and 1 indel) exclusive to brown morph *D. elegans* and *D. gunungcola*. It appears to be more diverged polymorphisms between species and similar shared polymorphisms between body color morphs to species comparisons. Therefore, both phylogenetic and polymorphism analyses do not support the evolutionary scenario of shared ancestral genetic variants.

## Discussion

### ebony: key driver of intraspecific and interspecific dimorphic pigmentation differences

Research on *Drosophila* pigmentation has revealed substantial variation in thorax, abdominal, and wing pigmentation shaped by selection pressures, geographic variables, and genetic factors. Notably, the sex chromosome and Muller element E (3R chromosome) harbor three key pigmentation genes, *yellow*, *tan*, and *ebony*, that play a central role in pigmentation divergence across *Drosophila* populations and species. Prior studies across *Drosophila* species have shown *ebony*’s key role in pigmentation spectrum (Massey & Wittkopp, 2016; Rebeiz et al., 2009; Takahashi et al., 2007; Wittkopp et al., 2009). Our findings, along with those of Massey et al. (2021), examined divergence between brown *D. elegans* and black *D. gunungcola*, highlighting *ebony* as a potential genetic hotspot underlying dimorphic pigmentation traits within and between species.

In *D. elegans*, dimorphism in thorax pigmentation appears to result primarily from *cis*-regulatory differences in *ebony*, rather than coding sequence mutations. The crucial role of *cis*-regulatory changes in the *ebony* locus has also been well-documented in several *Drosophila* species (Miyagi et al., 2015; Takahashi et al., 2007; Takahashi & Takano-Shimizu, 2011; Telonis-Scott et al., 2011). One of the most intriguing aspects of *ebony* is its pleiotropic nature, affecting not only pigmentation but also cuticular hydrocarbon, vision, and behavioral rhythmicity (Borycz et al., 2002; Massey et al., 2019; Takahashi, 2013). Evolution of the *cis-*regulatory region likely provides a more adaptable route in pleiotropic loci. In this case, the long 5’ intergenic region and intron 1 in the *ebony* locus offer opportunities to modulate the control for gene expression pattern and adaptive responses among multiple biological functions, thus making *ebony* an ideal target for selection.

In fact, the repetitive evolution has also been observed across various species, where the same sets of genes and pathways are repeatedly utilized within and between organisms. This process can lead to the emergence of novel traits or the development of similar traits across distinct species or populations over time. One of the well-known examples is the stickleback fish, where recurrent mutations of the *Eda* gene have independently led to the reduction of armor plates in multiple freshwater populations (Archambeault et al., 2020; Colosimo et al., 2005). This demonstrates how the same genetic changes can repeatedly drive similar adaptations in certain environments.

Our results suggest that repetitive evolution in dimorphic pigmentation differences within and between species may be associated with regulatory genetic changes across lineages. Similar phenotypes might evolve repeatedly in responding to parallel selective pressure in distinct populations or species. Further studies are required to explore how other genetic and regulatory pathways interact with *ebony*, which may uncover more profound insights into the mechanisms underlying such cases. Such research will broaden our understanding of evolutionary biology within *Drosophila* and shed light on the predictability of evolutionary changes across taxa.

### Repetitive evolution of pigmentation linked to independent regulatory changes

Repetitive evolution can arise through shared genetic polymorphism (collateral evolution), introgression, or independent genetic variation (parallel or convergent evolution), depending on whether pre-existing alleles or new mutations are involved in the process (Cerca, 2023; Maeso et al., 2012; Stern, 2013). The phylogenetic trees constructed from the 400-bp sequence in 5’ intergenic region (5’IR), coding sequence (CDS), and mitochondrial DNA (mtDNA), offer new insights into the evolutionary trajectories of dimorphic pigmentation within *D. elegans* species subgroup.

We use the mtDNA tree as a baseline to infer species-level divergence. The lack of distinct clades for brown and black morphs implies that there is no clear lineage sorting for pigmentation traits at the species level. However, the *ebony* CDS tree shows that black morphs nested within the brown morph clade, implying that black morphs might be a more recent adaptation or a variation that evolved from the brown morph *D. elegans* population. These contrasting patterns between mtDNA and *ebony* CDS may suggest that the traits are diversifying under selection pressures, while the overall population structure remains interconnected.

In contrast, the 5’IR tree shows two distinct clades of black and brown *D. elegans*, with *D. gunungcola* as a sister species. This phylogenetic pattern corresponds to ASE analysis, which indicates the link between *cis-*regulatory changes and the dimorphic pigmentation differences at both intra- and interspecific levels. Although this 400-bp region is relatively conserved between *D. elegans* and *D. gunungcola*, low sequence similarity of corresponding regions exists in other *Drosophila* species, indicating potential lineage specificity. Notably, we observed a relatively even distribution and number of shared genetic variants between black *D. elegans-D. gunungcola* contrast and brown *D. elegans-D. gunungcola* contrast. This balanced pattern does not support a scenario of shared ancestral variant or introgression, but instead suggests the involvement of independent evolutionary changes for the dimorphic pigmentation in the *D. elegans* species subgroup.

The pleiotropic effect of *ebony* in cuticular hydrocarbon (CHC) composition may also synergistically facilitate the evolutionary changes of *ebony* expression level in the epidermis. The CHC composition plays a significant role in desiccation resistance (Wang et al., 2022) and mate recognition (Chung & Carroll, 2015; Ferveur, 1997). The genetic ablation of *ebony* in *D. melanogaster, D. americana*, and *D. novamexicana* results in qualitative changes of CHC profiles with increased abundance of long chain CHCs (Lamb et al., 2020; Massey et al., 2019). In *D.* elegans, females of black and brown morphs exhibit significant differences in CHC profiles, potentially influencing male mating preference (Ishii et al., 2001). Thus, the interaction among desiccation resistance, mate recognition, and body-color related adaptation may maintain a strong selection imposing lower expression level of *ebony* in the epidermis of black morph *D. elegans* and *D. gunungcola.* Still, whether the same or overlapping regulatory elements influence the differences of body pigmentation and CHCs needs further investigations.

Our results provide a valuable baseline for further studies into the genetic and ecological underpinnings of pigmentation divergence. However, it remains unclear whether the divergent upstream region of *ebony* contributes specifically to pigmentation or also influences other traits, such as CHC composition. Moreover, the emergence of similar pigmentation in geographically distinct populations and species suggests the potential for repetitive evolutionary changes, although the underlying environmental or selective pressures require further investigations. Future research should aim to disentangle these complex relationships and identify the regulatory elements and environmental drivers that shape pigmentation and related traits in this system.

## Conclusion

Our findings suggest that pigmentation differences at both intra- and interspecific levels may have evolved independently through modifications in a lineage-specific sequence within the 5’ intergenic region of *ebony*. This highlights the predictability of adaptive evolution, where similar phenotypes repeatedly arise via modifications in specific regulatory regions, potentially driven by parallel selective pressures. Future research, including functional validation of this candidate region will be essential to confirm the regulatory role and clarify its contribution to dimorphic pigmentation differences within *D. elegans* species subgroup.

## Acknowledgments

We thank Jonathan H. Massey for assistance with library preparation and genome assembly for QTL mapping, and for helpful feedback on the early version of the manuscript. We are also grateful to Nia Kurniawan, Karuniawan Puji Wicaksono, and Hagus Tarno from the University of Brawijaya for their support during field sampling in Indonesia, including assistance with research permits and the material transfer agreement. This study was funded by National Science and Technology Council grant MOST107-2311-B-008-002 and MOST110-2311-B-008-002 to S.-D.Y. This study was also funded by National Institutes of Health Funding grant R35GM118073 to P.J.W. and National Science Foundation grant DBI 2209011 to E.X.B. The field trip for fly collection was funded by Ministry of Education grant to S.-D.Y.

## Data Accessibility Statement

All sequencing data have been deposited in National Center for Biotechnology Information (NCBI) under accession numbers listed in Table S2. All other relevant data, including QTL genotype and phenotype matrices, RT-qPCR results, and image files used for this study will be made publicly available via a suitable data repository upon acceptance.

## Benefits-Sharing Statement

Benefits from this research arise from the sharing of our data and findings through public databases, as described above, facilitating further research.

## Author contributions

S.-D.Y., B.-Y.L., and P.J.W. designed the project. A.S.F. wrote the manuscript with comments from all authors, analyzed the data, generated recombinants for QTL mapping, performed phenotyping, RT-qPCR, and ASE analyses. A.S.F. and S.-D.Y. collected wild flies. A.N. processed and phased long-read sequences to assemble haplotypes of the *ebony* region from wild-caught samples. K.-A.Y. extracted genomic DNA and prepared PCR amplicons for sequencing of the *ebony* region. E.X.B. performed HCR experiments and analyzed the resulting data. All authors read and approved the final manuscript.

## References

1. Andolfatto, P., Davison, D., Erezyilmaz, D., Hu, T. T., Mast, J., Sunayama-Morita, T., & Stern, D. L. (2011). Multiplexed shotgun genotyping for rapid and efficient genetic mapping. Genome Res, 21(4), 610–617. 10.1101/gr.115402.110

2. Archambeault, S. L., Bartschi, L. R., Merminod, A. D., & Peichel, C. L. (2020). Adaptation via pleiotropy and linkage: Association mapping reveals a complex genetic architecture within the stickleback Eda locus. Evol Lett, 4(4), 282–301. 10.1002/evl3.175

3. Bayala, E. X., Sinha, P., & Wittkopp, P. J. (2024). Protocol for dissecting Drosophila pupae and visualizing RNA expression using hybridization chain reaction. STAR Protoc, 5(4), 103456. 10.1016/j.xpro.2024.103456

4. Bohutinska, M., & Peichel, C. L. (2024). Divergence time shapes gene reuse during repeated adaptation. Trends in Ecology & Evolution, 39(4), 396–407. 10.1016/j.tree.2023.11.007

5. Borycz, J., Borycz, J. A., Loubani, M., & Meinertzhagen, I. A. (2002). tan and ebony genes regulate a novel pathway for transmitter metabolism at fly photoreceptor terminals. J Neurosci, 22(24), 10549–10557. 10.1523/JNEUROSCI.22-24-10549.2002

6. Brisson, J. A., De Toni, D. C., Duncan, I., & Templeton, A. R. (2005). Abdominal pigmentation variation in drosophila polymorpha: geographic variation in the trait, and underlying phylogeography. Evolution, 59(5), 1046–1059. https://www.ncbi.nlm.nih.gov/pubmed/16136804

7. Broman, K. W., Wu, H., Sen, S., & Churchill, G. A. (2003). R/qtl: QTL mapping in experimental crosses. Bioinformatics, 19(7), 889–890. 10.1093/bioinformatics/btg112

8. Cande, J., Andolfatto, P., Prud’homme, B., Stern, D. L., & Gompel, N. (2012). Evolution of multiple additive loci caused divergence between Drosophila yakuba and D. santomea in wing rowing during male courtship. PLoS One, 7(8), e43888. 10.1371/journal.pone.0043888

9. Carr, I. M., Robinson, J. I., Dimitriou, R., Markham, A. F., Morgan, A. W., & Bonthron, D. T. (2009). Inferring relative proportions of DNA variants from sequencing electropherograms. Bioinformatics, 25(24), 3244–3250. 10.1093/bioinformatics/btp583

10. Cerca, J. (2023). Understanding natural selection and similarity: Convergent, parallel and repeated evolution. Mol Ecol, 32(20), 5451–5462. 10.1111/mec.17132

11. Chung, H., & Carroll, S. B. (2015). Wax, sex and the origin of species: Dual roles of insect cuticular hydrocarbons in adaptation and mating. Bioessays, 37(7), 822–830. 10.1002/bies.201500014

12. Colosimo, P. F., Hosemann, K. E., Balabhadra, S., Villarreal, G., Jr., Dickson, M., Grimwood, J., Schmutz, J., Myers, R. M., Schluter, D., & Kingsley, D. M. (2005). Widespread parallel evolution in sticklebacks by repeated fixation of Ectodysplasin alleles. Science, 307(5717), 1928–1933. 10.1126/science.1107239

13. Courtier-Orgogozo, V., Arnoult, L., Prigent, S. R., Wiltgen, S., & Martin, A. (2020). Gephebase, a database of genotype-phenotype relationships for natural and domesticated variation in Eukaryotes. Nucleic Acids Res, 48(D1), D696–D703. 10.1093/nar/gkz796

14. Cuthill, I. C., Allen, W. L., Arbuckle, K., Caspers, B., Chaplin, G., Hauber, M. E., Hill, G. E., Jablonski, N. G., Jiggins, C. D., Kelber, A., Mappes, J., Marshall, J., Merrill, R., Osorio, D., Prum, R., Roberts, N. W., Roulin, A., Rowland, H. M., Sherratt, T. N., . . . Caro, T. (2017). The biology of color. Science, 357(6350). 10.1126/science.aan0221

15. Darriba, D., Taboada, G. L., Doallo, R., & Posada, D. (2012). jModelTest 2: more models, new heuristics and parallel computing. Nat Methods, 9(8), 772. 10.1038/nmeth.2109

16. Ferveur, J. F. (1997). The pheromonal role of cuticular hydrocarbons in Drosophila melanogaster. Bioessays, 19(4), 353–358. 10.1002/bies.950190413

17. Gibert, J. M., Mouchel-Vielh, E., De Castro, S., & Peronnet, F. (2016). Phenotypic Plasticity through Transcriptional Regulation of the Evolutionary Hotspot Gene tan in Drosophila melanogaster. PLoS Genet, 12(8), e1006218. 10.1371/journal.pgen.1006218

18. Haley, C. S., & Knott, S. A. (1992). A simple regression method for mapping quantitative trait loci in line crosses using flanking markers. Heredity (Edinb), 69(4), 315–324. 10.1038/hdy.1992.131

19. Hufnagel, D. E., Hufford, M. B., & Seetharam, A. S. (2020). SequelTools: a suite of tools for working with PacBio Sequel raw sequence data. BMC Bioinformatics, 21(1), 429. 10.1186/s12859-020-03751-8

20. Ishii, K., Hirai, Y., Katagiri, C., & Kimura, M. T. (2001). Sexual isolation and cuticular hydrocarbons in Drosophila elegans. Heredity (Edinb), 87(Pt 4), 392–399. 10.1046/j.1365-2540.2001.00864.x

21. Izore, T., Tailhades, J., Hansen, M. H., Kaczmarski, J. A., Jackson, C. J., & Cryle, M. J. (2019). Drosophila melanogaster nonribosomal peptide synthetase Ebony encodes an atypical condensation domain. Proc Natl Acad Sci U S A, 116(8), 2913–2918. 10.1073/pnas.1811194116

22. Johnson, W. C., Ordway, A. J., Watada, M., Pruitt, J. N., Williams, T. M., & Rebeiz, M. (2015). Genetic Changes to a Transcriptional Silencer Element Confers Phenotypic Diversity within and between Drosophila Species. PLoS Genet, 11(6), e1005279. 10.1371/journal.pgen.1005279

23. Kumar, S., Stecher, G., Li, M., Knyaz, C., & Tamura, K. (2018). MEGA X: Molecular Evolutionary Genetics Analysis across Computing Platforms. Mol Biol Evol, 35(6), 1547–1549. 10.1093/molbev/msy096

24. Kurtz, S., Phillippy, A., Delcher, A. L., Smoot, M., Shumway, M., Antonescu, C., & Salzberg, S. L. (2004). Versatile and open software for comparing large genomes. Genome Biol, 5(2), R12. 10.1186/gb-2004-5-2-r12

25. Lamb, A. M., Wang, Z., Simmer, P., Chung, H., & Wittkopp, P. J. (2020). ebony affects pigmentation divergence and cuticular hydrocarbons in Drosophila americana and D. novamexicana. Front Ecol Evol, 8. 10.3389/fevo.2020.00184

26. Maeso, I., Roy, S. W., & Irimia, M. (2012). Widespread recurrent evolution of genomic features. Genome Biol Evol, 4(4), 486–500. 10.1093/gbe/evs022

27. Martin, A., & Orgogozo, V. (2013). The Loci of repeated evolution: a catalog of genetic hotspots of phenotypic variation. Evolution, 67(5), 1235–1250. 10.1111/evo.12081

28. Massey, J. H., Akiyama, N., Bien, T., Dreisewerd, K., Wittkopp, P. J., Yew, J. Y., & Takahashi, A. (2019). Pleiotropic Effects of ebony and tan on Pigmentation and Cuticular Hydrocarbon Composition in Drosophila melanogaster. Front Physiol, 10, 518. 10.3389/fphys.2019.00518

29. Massey, J. H., Li, J., Stern, D. L., & Wittkopp, P. J. (2021). Distinct genetic architectures underlie divergent thorax, leg, and wing pigmentation between Drosophila elegans and D. gunungcola. Heredity (Edinb), 127(5), 467–474. 10.1038/s41437-021-00467-0

30. Massey, J. H., Rice, G. R., Firdaus, A. S., Chen, C. Y., Yeh, S. D., Stern, D. L., & Wittkopp, P. J. (2020). Co-evolving wing spots and mating displays are genetically separable traits in Drosophila. Evolution, 74(6), 1098–1111. 10.1111/evo.13990

31. Massey, J. H., & Wittkopp, P. J. (2016). The Genetic Basis of Pigmentation Differences Within and Between Drosophila Species. Curr Top Dev Biol, 119, 27–61. 10.1016/bs.ctdb.2016.03.004

32. Miyagi, R., Akiyama, N., Osada, N., & Takahashi, A. (2015). Complex patterns of cis-regulatory polymorphisms in ebony underlie standing pigmentation variation in Drosophila melanogaster. Mol Ecol, 24(23), 5829–5841. 10.1111/mec.13432

33. Parkash, R., Sharma, V., & Kalra, B. (2008). Climatic adaptations of body melanisation in Drosophila melanogaster from Western Himalayas. Fly (Austin), 2(3), 111–117. 10.4161/fly.6351

34. Pfaffl, M. W. (2001). A new mathematical model for relative quantification in real-time RT-PCR. Nucleic Acids Res, 29(9), e45. 10.1093/nar/29.9.e45

35. Pool, J. E., & Aquadro, C. F. (2007). The genetic basis of adaptive pigmentation variation in Drosophila melanogaster. Mol Ecol, 16(14), 2844–2851. 10.1111/j.1365-294X.2007.03324.x

36. R Core Team. (2024). R: A Language and Environment for Statistical Computing. R Foundation for Statistical Computing. https://www.R-project.org/

37. Rebeiz, M., Pool, J. E., Kassner, V. A., Aquadro, C. F., & Carroll, S. B. (2009). Stepwise modification of a modular enhancer underlies adaptation in a Drosophila population. Science, 326(5960), 1663–1667. 10.1126/science.1178357

38. Richardt, A., Kemme, T., Wagner, S., Schwarzer, D., Marahiel, M. A., & Hovemann, B. T. (2003). Ebony, a novel nonribosomal peptide synthetase for beta-alanine conjugation with biogenic amines in Drosophila. J Biol Chem, 278(42), 41160–41166. 10.1074/jbc.M304303200

39. Ronquist, F., Teslenko, M., van der Mark, P., Ayres, D. L., Darling, A., Hohna, S., Larget, B., Liu, L., Suchard, M. A., & Huelsenbeck, J. P. (2012). MrBayes 3.2: efficient Bayesian phylogenetic inference and model choice across a large model space. Syst Biol, 61(3), 539–542. 10.1093/sysbio/sys029

40. Rozas, J., Ferrer-Mata, A., Sanchez-DelBarrio, J. C., Guirao-Rico, S., Librado, P., Ramos-Onsins, S. E., & Sanchez-Gracia, A. (2017). DnaSP 6: DNA Sequence Polymorphism Analysis of Large Data Sets. Mol Biol Evol, 34(12), 3299–3302. 10.1093/molbev/msx248

41. Stern, D. L. (2013). The genetic causes of convergent evolution. Nat Rev Genet, 14(11), 751–764. 10.1038/nrg3483

42. Takahashi, A. (2013). Pigmentation and behavior: potential association through pleiotropic genes in Drosophila. Genes Genet Syst, 88(3), 165–174. 10.1266/ggs.88.165

43. Takahashi, A., Takahashi, K., Ueda, R., & Takano-Shimizu, T. (2007). Natural variation of ebony gene controlling thoracic pigmentation in Drosophila melanogaster. Genetics, 177(2), 1233–1237. 10.1534/genetics.107.075283

44. Takahashi, A., & Takano-Shimizu, T. (2011). Divergent enhancer haplotype of ebony on inversion In(3R)Payne associated with pigmentation variation in a tropical population of Drosophila melanogaster. Mol Ecol, 20(20), 4277–4287. 10.1111/j.1365-294X.2011.05260.x

45. Telonis-Scott, M., Hoffmann, A. A., & Sgro, C. M. (2011). The molecular genetics of clinal variation: a case study of ebony and thoracic trident pigmentation in Drosophila melanogaster from eastern Australia. Mol Ecol, 20(10), 2100–2110. 10.1111/j.1365-294X.2011.05089.x

46. Wang, Z., Receveur, J. P., Pu, J., Cong, H., Richards, C., Liang, M., & Chung, H. (2022). Desiccation resistance differences in Drosophila species can be largely explained by variations in cuticular hydrocarbons. Elife, 11. 10.7554/eLife.80859

47. Wittkopp, P. J., Stewart, E. E., Arnold, L. L., Neidert, A. H., Haerum, B. K., Thompson, E. M., Akhras, S., Smith-Winberry, G., & Shefner, L. (2009). Intraspecific polymorphism to interspecific divergence: genetics of pigmentation in Drosophila. Science, 326(5952), 540–544. 10.1126/science.1176980

48. Wittkopp, P. J., True, J. R., & Carroll, S. B. (2002). Reciprocal functions of the Drosophila yellow and ebony proteins in the development and evolution of pigment patterns. Development, 129(8), 1849–1858. 10.1242/dev.129.8.1849

